# A ubiquitous and diverse methanogenic community drives microbial methane cycling in eutrophic coastal sediments

**DOI:** 10.1101/2024.10.04.616656

**Authors:** A.J. Wallenius, J. Venetz, O.M. Zygadlowska, W.K. Lenstra, N.A.G.M. van Helmond, P. Dalcin Martins, C.P. Slomp, M.S.M. Jetten

**Affiliations:** Department of Microbiology, Radboud Institute for Biological and Environmental Sciences, Radboud University, Nijmegen, the Netherlands; Department of Earth Sciences - Geochemistry, Utrecht University, Utrecht, the Netherlands; Institute for Biodiversity and Ecosystem Dynamics, University of Amsterdam, The Netherlands

**Keywords:** Methanogens, ANME, coastal sediment, microbial methane cycle

## Abstract

Coastal areas are responsible for over 75% of global marine methane emissions and this proportion is predicted to grow due to an increase in anthropogenically induced eutrophication and deoxygenation. Prolonged periods of low oxygen and high organic matter input have been put forward to cause an imbalanced microbial methane cycle, as methane oxidation cannot keep up with methane production. However, it is still unclear what factors affect each process and which microorganisms are responsible. Here we show that methanogenic processes dominate microbial methane cycling in the anoxic sediments of marine Lake Grevelingen (NL) after summer stratification with bottom water anoxia. We observed a shallow and narrow sulfate-methane transition zone between 5 and 15 cm depth, with high methane concentrations (> 5 mM) below this zone. Methanogenesis was dominant over methanotrophy in all investigated layers as active methanogenesis potential was detected down to 60 cm below sea floor, but methane oxidation was only observed in a narrow section of the sulfate-methane transition zone. Based on amplicon sequencing and sediment incubations, we uncovered a metabolically and phylogenetically diverse methanogenic community with distinct niche separation in different sediment layers. ANME archaea and their putative syntrophic sulfate-reducing bacteria were restricted to a narrow zone and were co-occurring with the detected methane oxidation activity. Our results suggest that eutrophication and deoxygenation will further contribute to rising methane emissions in coastal areas, as the microbial methane cycle will be tilted towards increased methanogenesis while the efficiency of the microbial methane filter is expected to decline.

## Introduction

Methanogenic archaea in marine sediments produce methane (Reeburgh, 2007; Saunois et al., 2020), a greenhouse gas 86 times more potent than CO_2_ on a 20-year time scale. Atmospheric methane concentrations are rising sharply with emissions from both anthropogenic and biogenic sources contributing significantly (Saunois et al., 2020). Most of the methane produced in deep marine sediments does not reach the atmosphere, as anaerobic methanotrophic archaea (ANME), in consortium with sulfate-reducing bacteria (SRB), are estimated to consume over 90% of the biologically produced methane (Knittel and Boetius, 2009). This sulfate-dependent anaerobic oxidation of methane (S-AOM) takes place in the so-called sulfate-methane transition zone (SMTZ) located above the methanogenic sediment layer.

In coastal and shelf areas, the SMTZ is often found to be narrower and at a shallower depth than in the deeper parts of the oceans (Egger et al., 2018), which might affect its methane removal capacity, thus allowing methane to bypass the SMTZ and to escape to the atmosphere (Żygadłowska et al., 2024a). The drivers of the imbalance between methane production and oxidation in coastal systems are not well-explored, but anthropogenic activity-driven eutrophication and resulting high organic matter (OM) input and increased hypoxia often favour methanogenesis over methanotrophs (Wallenius et al., 2021).

In eutrophic systems, high sedimentation rates can lead to a shoaling of the SMTZ, and consequently, bring the methanogenic zone closer to the sediment-water interface (Egger et al., 2016; Żygadłowska et al., 2024a). Increased hypoxia in coastal bottom waters will also shift the redox zones upwards as oxygen and nitrate are no longer available as an electron acceptor in the top sediments. Subsequently, labile organic matter will be buried in higher quantities below the SMTZ, providing more substrate for fermenters and ultimately methanogens. Furthermore, more available substrates lead to less competition and enable methanogens to thrive even in and above the SMTZ (Xiao et al., 2017; Beulig et al., 2019; Coon et al., 2023). Thus, although coastal areas account for more than 75 % of total marine methane emissions (while only covering 2% of the marine surface area) these emissions are expected to increase with ongoing eutrophication and deoxygenation (Rosentreter et al., 2021).

In marine sediments, H_2_/CO_2_, and formate are the most prevalent methanogenic substrates (hydrogenotrophic methanogenesis), followed by acetate (acetoclastic methanogensis; Liu and Whitman, 2008). Methylated substrates, such as methanol and methylated amines can also be converted to methane (methylotrophic methanogenesis), with or without H_2_ as a reductant, but their significance in marine environments has only gained more attention in recent years (Fischer et al., 2021; Tsola et al., 2024). As acetate and hydrogen can also be consumed efficiently by SRB, methylated compounds were long assumed to be “non-competitive” substrates for methanogens. However, with the isolation of several methylotrophic SRB this assumption should probably be revised (Sousa et al., 2018). Some methylated substrates such as trimethylamine and dimethylsulfide (DMS) are abundant in vegetated and hypersaline sediments and can be the main contributors to methane emissions. Also, many novel archaeal lineages encode the genes for methylotrophic methanogenesis (Evans et al., 2015; Vanwonterghem et al., 2016; McKay et al., 2019), and the first cultured representatives have recently been isolated (Kohtz et al., 2024; Krukenberg et al., 2024; Wu et al., 2024), expanding the occurrence and significance of this pathway.

Marine sediments generally harbour a diverse methanogenic community that can change rapidly with depth due to changing porewater chemistry and available substrates (Borrel et al., 2012; Zhang et al., 2020). Most methanogens only encode the machinery for one methanogenic pathway, and members of the same order commonly have a very similar metabolism such as hydrogenotrophic Methanomicrobiales or Methanobacteriales (Liu and Whitman, 2008; Evans et al., 2019). The order Methanosarcinales harbors the most diverse metabolisms including methylotrophic species such as the common marine sediment inhabitant *Methanolobus*, the metabolically versatile *Methanosarcina*, and the strictly acetoclastic *Methanosaeta* (*Methanothrix*). In addition, marine methane-oxidizing ANME archaea are also members of Methanosarcinales, but they belong to several different families (Chadwick et al., 2022).

Coastal systems often have lower salinity and thus less sulfate available for S-AOM. Thus, with the expanding methanogenic zone and increased methane production in eutrophic coastal sediments, the methanotrophic S-AOM community will be confronted with a methane surplus challenge. As ANME archaea are notoriously slow-growing (Dale et al., 2008; Knittel et al., 2018; Lenstra et al., 2023), they might not be able to establish a stable methane oxidizing zone (Egger et al., 2016). In the absence of sulfate, some ANME clades might be able to couple methane oxidation to metal oxides (metal-AOM) or nitrate (N-AOM) reduction (Wallenius et al., 2021), but the significance of these processes to the overall methane removal potential in coastal systems is not well quantified yet.

When methane production exceeds methanotrophy in coastal sediments, the methane cycle might become out of balance, resulting in increased benthic methane flux and emissions to the atmosphere (Venetz et al., 2023; Żygadłowska et al., 2023). With intensifying anthropogenic activity in coastal areas, eutrophication and hypoxia are predicted to further expand (Breitburg et al., 2018; Sinha et al., 2017). This may lead to even higher methane emissions from coastal sediments, making it urgent to understand the factors influencing the methane cycling processes and microorganisms.

Here, we investigated the methane cycling potential and the metabolic pathways in anoxic sediments of eutrophic coastal marine Lake Grevelingen (the Netherlands). We aimed to answer the following questions: 1) What are the rates of methanogenesis and methanotrophy across the sediment, 2) What are the key microorganisms driving methane cycling in the different redox zones; and 3) Which metabolic pathways are used for methane production and oxidation. These questions were addressed by a complementary set of experiments involving sediment incubation experiments to determine the rates of the various processes, DNA extractions together with 16S rRNA amplicon sequencing to assess the diversity and the potential methane cycle processes.

## Materials & methods

### Study site and sampling

The Scharendijke basin (45 m; 51.742°N; 3.849°E) is the deepest part of Lake Grevelingen, located in the southwestern part of the Netherlands. The lake used to be an estuary until it was dammed, and partly reconnected to the North Sea in 1978 (Bannink et al., 1984), and presently has a salinity of 29-32. The sedimentation rate in the basin is high (up to 20 cm yr^-1^). The water column is subject to seasonal stratification, resulting in euxinic bottom waters in the summer months (Egger et al., 2016; Hagens et al., 2015; Żygadłowska et al., 2023). Sediment cores were collected in September 2020 during a sampling campaign on board R/V Navicula with a UWITEC gravity corer and sliced anoxically for incubation experiments or subsampled and immediately frozen at -80°C for DNA analysis. Sliced cores were stored anoxically, in gas tight aluminium bags flushed with N_2_ gas, in the dark at 4°C until the start of incubation experiments within 2-4 months after sampling Porewater samples for CH_4_, SO_4_^2-^, H_2_S, alkalinity and δ^13^C-CH_4_ were collected directly after core collection and analysed as described in Żygadłowska et al.(2023).

### Methanogenic incubations

Sediments from selected depths were diluted with anoxic sulfate-free artificial seawater (ASW) medium in 1:1 ratio in an anaerobic chamber and carefully homogenized. Then, 10 g of sediment slurry was transferred into 60 ml serum bottles which were immediately stoppered and capped. The artificial sea water medium with a salinity of 32 was composed of NaCl (26 g), MgCl_2_·6H_2_O (7.48 g), CaCl_2_·2H_2_O (1.45 g), KCl (0.6 g) and 20 mL of NaHCO_3_ solution (1M) per liter of demineralized water. The pH was adjusted to 7.8. To remove any trace gases, the samples were flushed with argon (Ar), leaving 1.5 bar pressure, and the bottles were preincubated in the dark at 4ºC for three weeks. Before substrate addition, the headspace was flushed with Ar to remove any methane produced during pre-incubation and replaced with 99.5% Ar and 0.5% CO_2_. For each depth, a duplicate sample was prepared without any added substrates to observe the endogenous methane production. For three selected depths (5-15 cm, 25-35 cm, 45-55 cm) the bottles were amended with acetate (5 mM), methanol (2 mM), methanol and H_2_ (2 mM; 8 mM) or CO_2_ and H_2_ (2 mM; 8 mM). Each condition was prepared in triplicate. The bottles were incubated at room temperature in the dark while gently shaking. After 33 days of incubation, when all substrates appeared to be consumed, a second dose of acetate (10 mM), methanol (6 mM), methanol and H_2_ (24 mM) or CO_2_ (6 mM) and H_2_ (24 mM) was added to the bottles, 2-3 fold more in order to maintain methanogenesis activity for longer period. Methane and hydrogen concentrations in the headspace were measured with a HP5890 gas chromatograph (Agilent Technologies, Santa Clara, CA, USA) equipped with a Porapak Q column and a thermal conductivity detector. Methane production rates were calculated for the first three days of incubations for all samples except for acetate, where substrate consumption only started after a two-week lag phase and thus rates were calculated from days 17-24 when the methane increase was linear.

### Methanotrophic incubations

The methane oxidation potential was determined for five selected depths. Ten grams of sediment was diluted in a 1:5 ratio with sulfate-free ASW in sterile 120 ml serum bottles, stoppered and capped. The headspace was replaced with 100% methane with 1.5 bar overpressure and the samples were pre-incubated for 3 weeks in the dark at 4°C to activate methane oxidizing microorganisms. To start the incubation, headspace was thoroughly flushed with Ar to remove all methane and amended with 20% ^13^C-CH_4_, 1% N_2_ and 1% CO_2_. The samples were amended with 5 mM NaSO_4_^2^ as electron acceptor for methane oxidation, or only ^13^C-CH_4_ as a control and incubated while gently shaking in the dark at room temperature. The ratio of ^13/12^C-CO_2_ was determined with a GC-MS (Agilent 5975C inert MSD).

### DNA isolation

The in-situ samples were defrosted and homogenized, and 0.2 g of the sediment slurry was bead-beaten on a TissueLyser LT (Qiagen, Venlo, Netherlands) for 10 min at 50 Hz in a PowerBead tube from the DNeasy PowerSoil DNA isolation kit (Qiagen) before isolating the DNA according to manufacturer’s instructions. The quantity and quality of isolated DNA eluted in sterile MilliQ water were assessed by NanoDrop 1000 (Thermo Fischer Scientific, Bremen, Germany), Qubit® 2.0 (Invitrogen, USA) and on 1% agarose gel. After isolation, DNA was immediately frozen at −20°C until further analysis.

### 16S rRNA gene amplicon sequencing and analysis

The amplification of the total archaeal 16S rRNA genes was performed with primers Arch349F (5′-GYGCASCAGKCGMGAAW-3′) and Arch806R (5′-GGACTACVSGGGTATCTAAT-3′; (Takai and Horikoshi, 2000). The primers used for bacterial 16S rRNA gene amplification were Bac341F (5′-CCTACGGGNGGCWGCAG-3′; Herlemann et al., 2011) and Bac806R (5′-GGACTACHVGGGTWTCTAAT-3′; Caporaso et al., 2012). 16S rRNA amplicon sequencing was performed on the Illumina MiSeq Next Generation Sequencing platform by Macrogen (Seoul, South Korea) using Herculase II Fusion DNA Polymerase Nextera XT Index Kit V2, yielding 2×300bp paired-end reads. The quality of raw reads was checked using FastQC (v0.11.5;(Andrew, 2010). The paired-end reads were trimmed with Cutadapt (v1.18; Martin, 2011) to remove adapters. The data were further processed using the DADA2 pipeline (v1.8; Callahan et al., 2016) in RStudio to cluster the reads into amplicon sequence variants (ASVs), chimera removal and taxonomic classification using SILVA 16S rRNA database (v138.1; Quast et al., 2013). Microbial community data analysis was performed using the package phyloseq (v1.36.0; McMurdie and Holmes, 2013) and visualized using ggplot2 (v3.3.5; Wickham, 2016). The raw sequence reads can be accessed with BioProject accession PRJNA1167897.

## Results

### Pore water profile and methanogenesis rates

The Scharendijke basin sediments in Lake Grevelingen are characterized by a high sedimentation rate and OM input, which is reflected by the shallow SMTZ at approximately 5-15 cm below the sediment-water interface (Fig. 1a, b). In the top 5 cm, sulfate concentrations decrease rapidly from 25 to 3 mM as sulfate acts as the primary electron acceptor for OM degradation by SRB, which results in high sulfide concentrations in the porewater in the same zone, peaking at 5 cm depth (up to 5 mM, Fig. 1c). The upward diffusing methane in the SMTZ was depleted in ^13^C, indicating S-AOM probably does not contribute much to the sediment methane signature. (Fig. 1d). Methane concentrations remain high (4-6 mM) down to 60 cm depth indicating vigorous methanogenesis and accumulation of methane throughout the sediment. The steep increase in alkalinity in the top 5 cm supports high OM degradation rates, and the steady downward increase suggests continued mineralization at greater depth although at lower rates, which could be advantageous to methanogens competing with SRB for substrates (Fig. 1e). Surprisingly, endogenous methanogenesis was observed at all incubated depths from 0-5 cm to 45-55 cm, albeit at a slow rate (0.1 ± 0.02 µmol methane g_dw_ ^-1^ day^-1^) in the top 5 cm (, Fig. 1f). Also, unexpectedly for marine sediments, the highest methanogenesis rate (1.7 ± 0.01 µmol methane g_dw_^-1^ day^-1^) was measured in the SMTZ at depths between 5-10 cm where the sulfate concentration was still above 2 mM. Below the SMTZ, the methane production rate remained stable at about 1.2 µmol methane g _dw_^-1^ day^-1^.

**Fig. 1.**
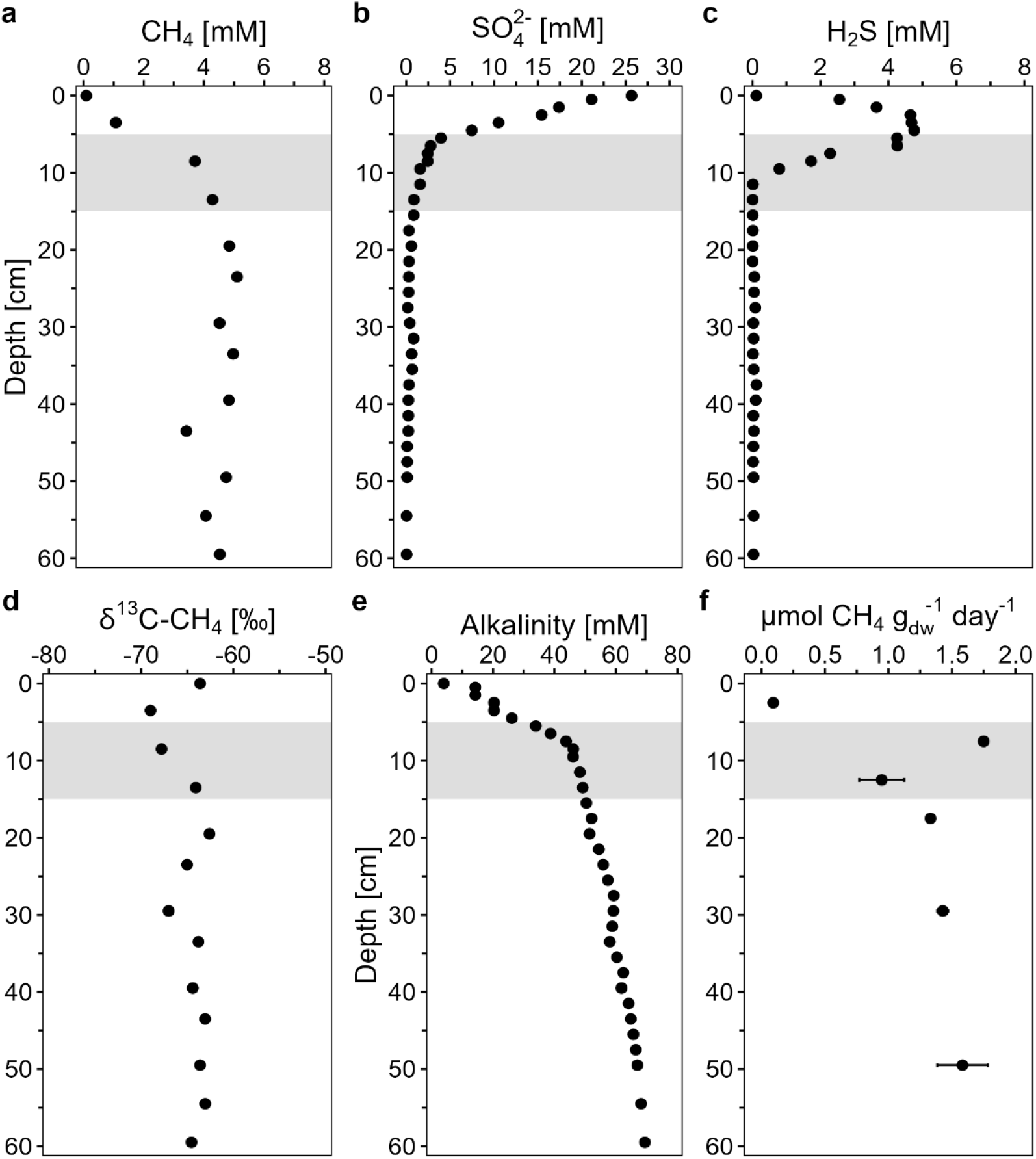
Porewater depth profile of a) methane, b) sulfate, c) sulfide, d) alkalinity, e) δ^13^C-CH_4_, and f) endogenous methanogenesis rates (n=2). The gray shaded area indicates the SMTZ. Full porewater and solid phase profiles can be seen in Żygadłowska et al., 2023.

### High methanogenesis potential across sediment and substrates

At all depths, all added methanogenic substrates initiated methane production well above the control levels (Fig. 2). From all the substrates, the highest methanogenesis rates were measured in the 5-15 cm incubations with methanol and H_2_ as substrates, with a maximum of 8.8 µmol CH_4_ g_dw_ ^-1^ day^-1^. Acetoclastic, hydrogenotrophic and methylotrophic (without H_2_) methanogenesis rates were similar (∼3.7 - 4.7 µmol CH_4_ g_dw_ ^-1^ day^-1^) across the various sediment intervals, except for the acetate amended bottles from the 5-15 cm layer where methane production was fastest (7.2 µmol CH_4_ g_dw_ ^-1^ day^-1^). Here, CO_2_/H_2_ amended sediment had the slowest methane production rate, suggesting that other methanogenic pathways are more active in the SMTZ. Also, at the deepest depth (45-55 cm) the measured methane production rates were comparable to the higher depths, indicating that there is still a substantial methanogenic community at this depth that can be easily activated. Overall, all depths showed a high potential for methanogenesis from all the main methanogenic substrates. When the incubations were provided with a second dose of substrate, the adapted methanogenic community showed an even higher maximum methane production rate of 30 µmol CH_4_ g_dw_ ^-1^ day^-1^ with methanol/H_2_ (Fig.S1a).

**Fig. 2.**
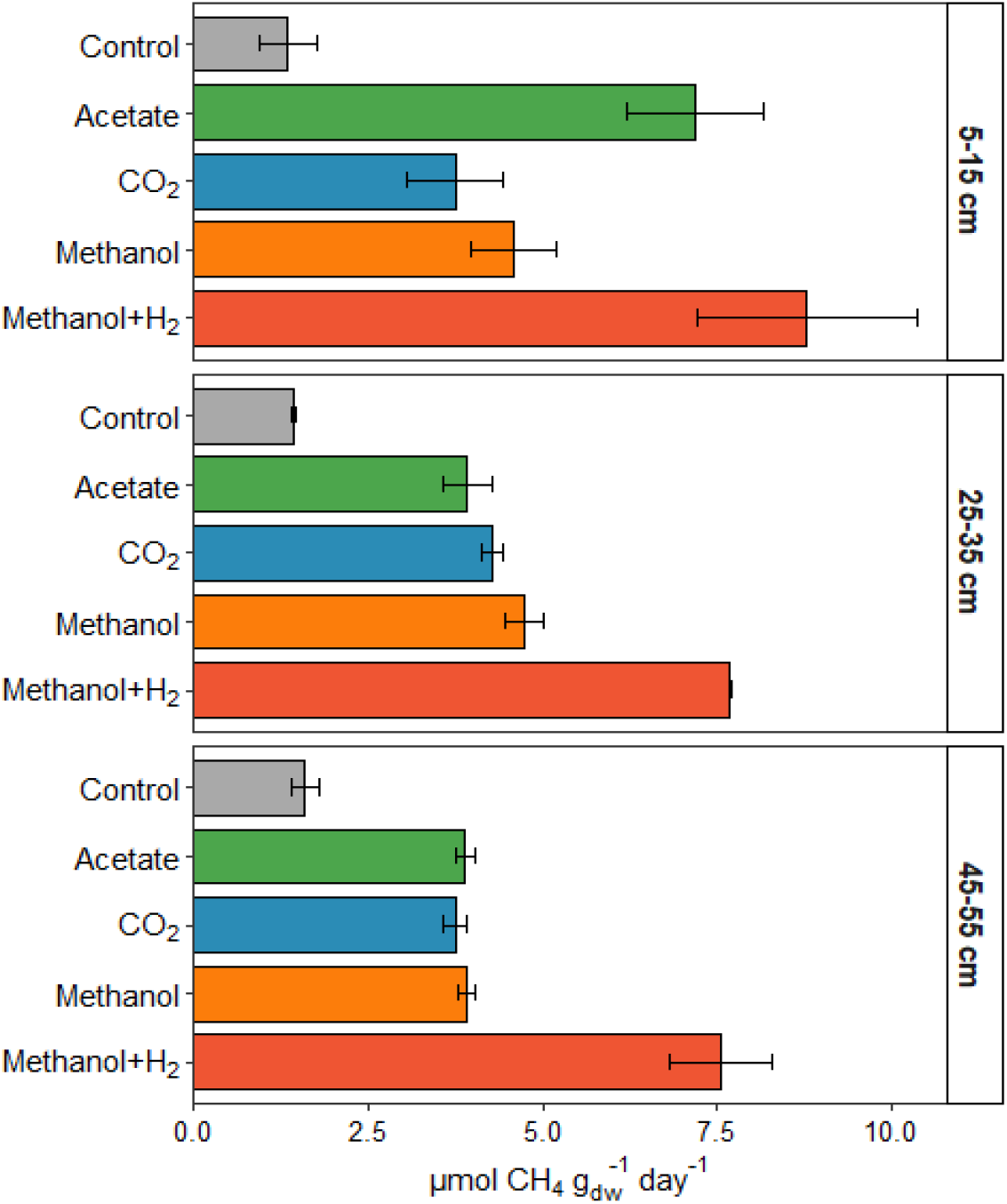
Potential methane production rates from different methanogenic substrates in Lake Grevelingen sediment from 3 different depths. Error bars represent variation between duplicates (control) or triplicates.

### Methane oxidation detected in narrow zone below SMTZ

In addition to the methane production potential, we investigated the change in anaerobic methane oxidation potential with depth. The increase in the ratio of ^13/12^C-CO_2_ is the result of oxidation of ^13^C-CH_4_ and is therefore taken as an indication for S-AOM. For the five tested depths above, at and below the SMTZ, active methane oxidation was only observed at 10-15 cm and 15-20 cm depth (Fig. 3a), with the highest increase in ^13^C-CO_2_ at 15-20 cm, directly below the SMTZ (Fig. 1a). The ^13/12^C ratio started to increase already after 1 day, suggesting an active S-AOM community in this narrow zone around the SMTZ. In the control samples without added electron acceptors, the ^13/12^C ratio increased only slightly in all depths, suggesting some residual electron acceptor and endogenous AOM activity at all depths, albeit at much lower levels than in the sulfate-amended samples (Fig. 3b).

**Fig. 3.**
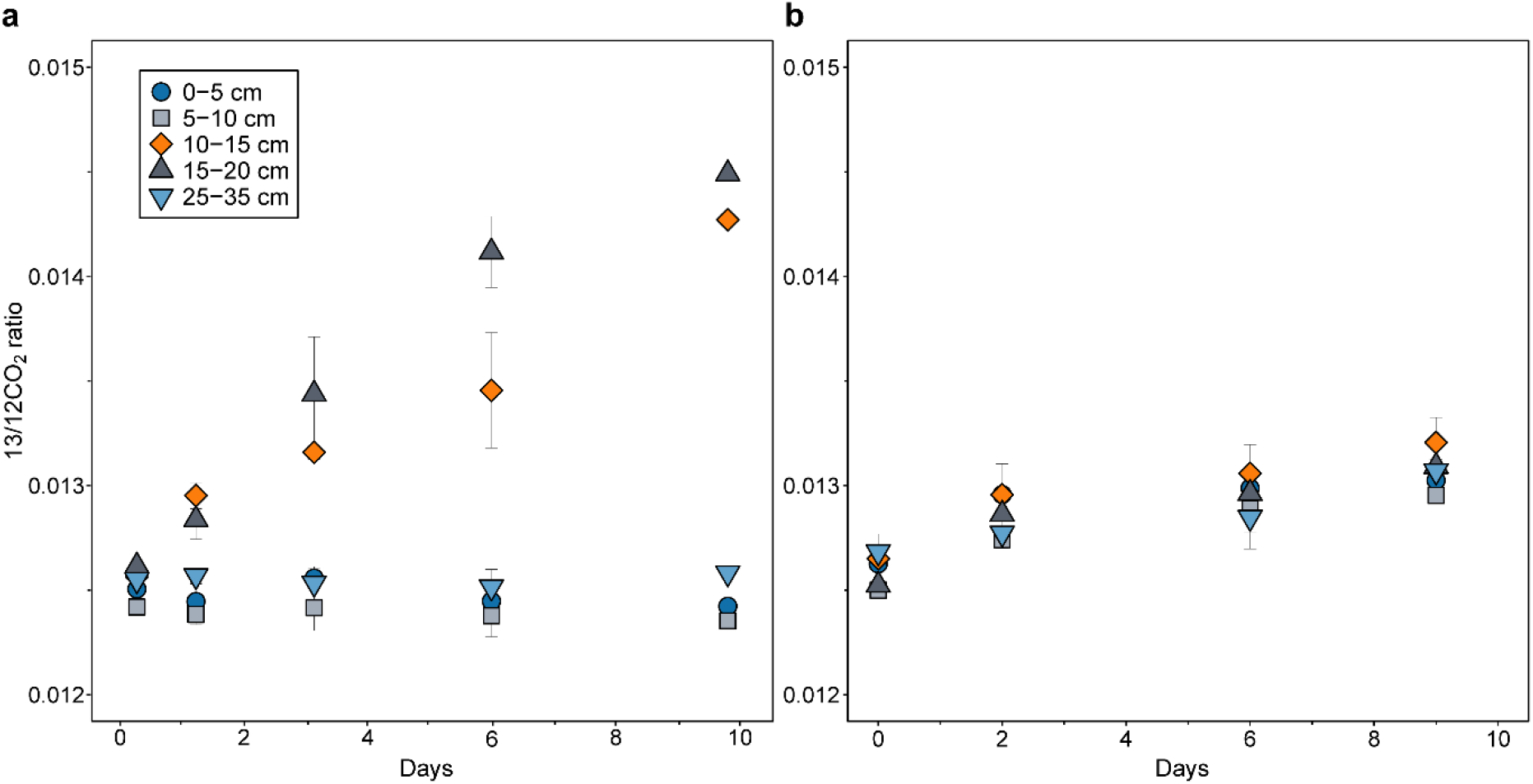
Potential S-AOM rates for 5 sediment depth intervals. The increase in the ratio of ^13/12^C-CO_2_ in the headspace gas phase in batch incubations amended with a) ^13^C-CH_4_ and sulfate, or b) ^13^C-CH_4_ only.

### Methanogens and ANME archaea inhabit specific niches in the sediment

Methanogenic archaea were found in high abundance across the top 60 cm of the sediment, and the diversity of methanogenic taxa was high (Fig. 4). Unlike in deep marine sediments, methanogenic archaea were detected in high abundance at and above the SMTZ, indicating that they can compete with SRB and other bacteria and archaea involved in OM degradation. Methylotrophic *Methanolobus* and *Methanococcoides* were the most abundant methanogens in the top 10 cm and amounted to 17% of the recovered archaeal reads, suggesting that the methane produced in the top layer originates from methylated substrates. However, hydrogenotrophic methanogens from the *Methanomicrobiaceae* family were also prominent in the top layers. The SMTZ did not harbour an overall high abundance of methanogens, but methylotophic genera were more abundant in the top part of the SMTZ with high sulfide, whereas other genera such as Methanosaeta and Methanogenium were more abundant around 10 cm when most sulfide was consumed. Below the SMTZ, methanogen abundance increased rapidly from 20% to up to 90% of archaeal reads at 40 cm, with hydrogenotrophic methanogens being the most dominant group. Strictly acetoclastic *Methanosaeta* were commonly found below 30 cm and increased in abundance when *Methanosarcina* and *Methanogenium* were decreasing, indicating differences in organic matter composition and available substrates.

**Fig. 4.**
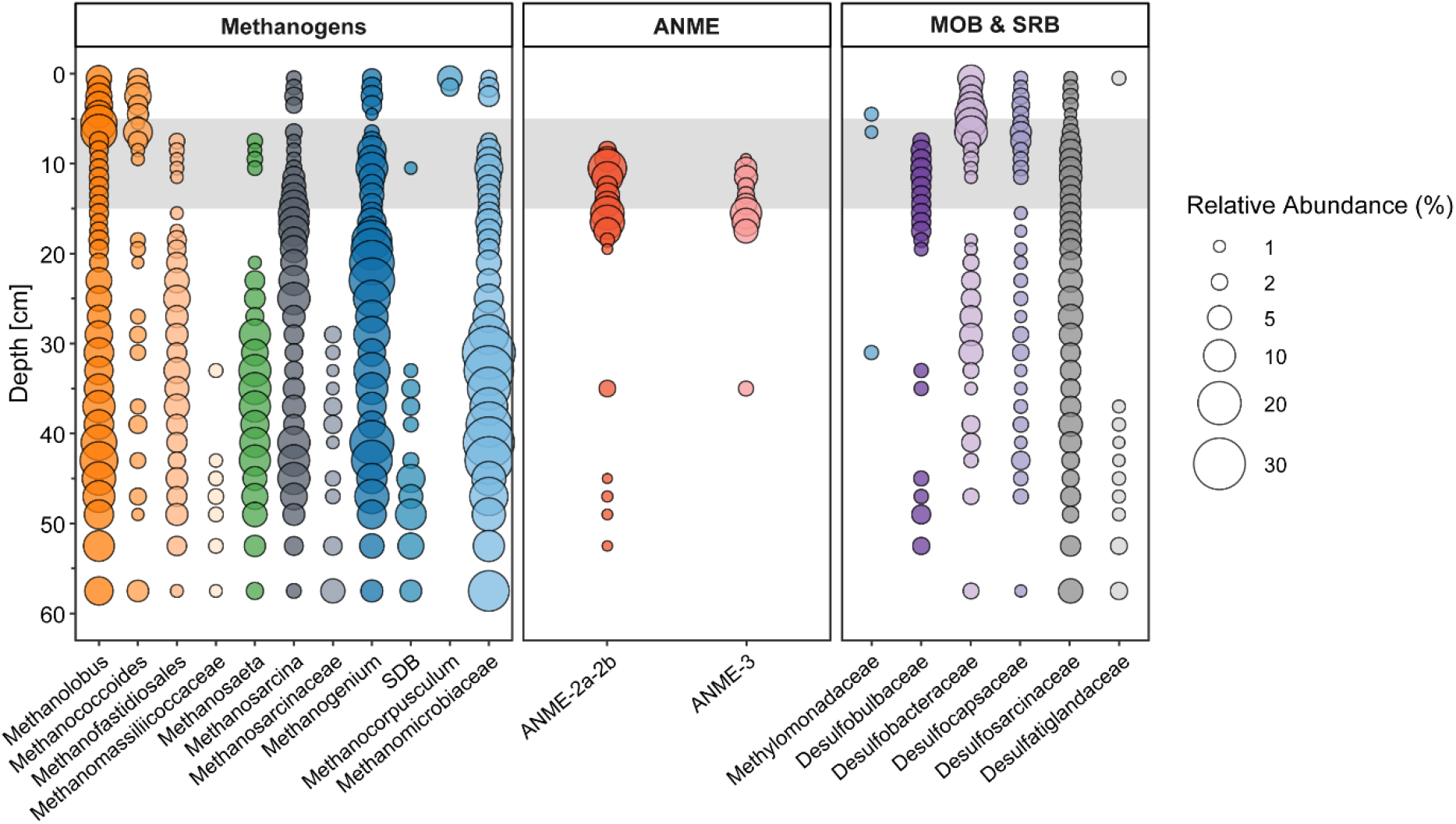
The relative abundance of methane-cycling archaea and methanotrophic and sulfate-reducing bacteria. The grey box represents the SMTZ. Methanogens are listed and grouped by the potential metabolic pathway in different colours; methylotrophic (orange); acetoclastic (green); flexible (grey); hydrogenotrophic (blue). MOB = Methane oxidizing bacteria (blue); SRB; sulfate reducing bacteria (purple/grey). Only taxa with >1% relative abundance in at least two depths are included.

Representative taxa of all main methanogenic guilds and pathways were ubiquitously present in the sediment, thus making it challenging to suggest a dominant methane source, but rather, methanogenesis in Lake Grevelingen sediments seems to proceed via multiple metabolic pathways conducted by a very diverse methanogenic community.

In contrast to the methanogen abundance, but in line with the S-AOM activity test, methanotrophic ANME were only detected in a narrow zone at and below the SMTZ. ANME-2a/2b accounted for up to 15% of the archaeal community at 10-11 cm, and ANME-3 10% at 15-16 cm. Each clade appeared to have a specific niche at the SMTZ, with ANME-2a/2b dominating at higher sulfate concentrations, and ANME-3 when sulfate was below 0.1 mM. The distribution of ANMEs is supported by the high abundance of *Desulfobulbaceae* reads in the SMTZ, a putative bacterial partner in S-AOM. Other SRB families did not seem to be dependent on ANME occurrence, as *Desulfobacteraceae* inhabited mostly the top sediments above the SMTZ, and *Desulfosarcinaceae* were more prominent across the sediment profile. However, on genus level, SEEP-SRB1 group was more abundant in the SMTZ than at other depths, suggesting a relationship with ANME (Fig. S2). In some samples ANME reads were detected below 35 cm, but in < 2% abundance, likely representing buried cells.

Despite the high methanogen abundance, high methane production potential and high sulfate concentration in the top 5 cm, anaerobic or aerobic methanotrophs were not detected in this zone. Bacterial *Methylomonadaceae* reads were detected around 5 cm depth, where they likely represent buried cells from the water column MOB community (Venetz et al., 2023). Thus, the methanogenic archaea seem to be dominant over methanotrophs throughout the top 60 cm of Lake Grevelingen sediments.

## Discussion

Coastal ecosystems are increasingly under pressure through eutrophication, warming and deoxygenation and may become bigger hotspots for methane. Eutrophication, a rapid rate of sedimentation, bottom water hypoxia and high methane diffusive and ebullitive fluxes have been frequently reported for marine Lake Grevelingen (Hagens et al., 2015; Egger et al., 2016; Sulu-Gambari et al., 2017; Żygadłowska et al., 2023). These previous studies on microbial methane cycling have focused on methane oxidation in the anaerobic sediment or in the stratified water column (Bhattarai et al., 2017, 2018; Venetz et al., 2023, 2024), but the methanogenic community and pathways have not been characterized before. Here, we investigated the overall microbial methane cycling in the sediment, potential production and removal pathways and key microbes involved, with an array of complementary methods such as biogeochemical profiles, activity tests and 16S rRNA profiles.

### Shallow SMTZ results in narrow redox zonation

At the end of summer after several months of water column stratification, the SMTZ at Scharendijke basin was very narrow and occurred close to the sediment-water interface at 5-15 cm (Fig 1). The SMTZ in coastal sediments is often found to be both much narrower and shallower in comparison to that in deep marine sediments, due to higher sedimentation rates (Egger et al., 2018). This can result in more methane escaping into the water column as the methanogenic zone is wider and more labile OM is buried below the SMTZ as a result of incomplete mineralization. The methane oxidation potential is also compromised, as ANME archaea cannot remove all the methane in the narrow niche. Żygadłowska et al. (2023) reported high methane concentrations in the bottom waters of Scharendijke after summer stratification and no noticeable signal for methane oxidation in the sediment based on the isotopic signature (Fig. 1d). The unusually high methane production rates and low values of δ^13^C-CH_4_ within the SMTZ confirm that methanogenesis outpaces methanotrophy in these sediments.

### Methanogenesis potential and methanogenic diversity are high throughout the sediment

In agreement with the high methane concentrations and the broad methanic zone in Lake Grevelingen, endogenous methanogenesis occurs at high rates (max. 1.7 µmol CH_4_ g _dw_^-1^ day ^-1^) across the top 60 cm of Scharendijke sediment, which is high for a coastal marine environment (Krüger et al., 2005; Chuang et al., 2016). In marine sediments, the methanic zone is usually located below the zone where sulfate is depleted where CO_2_ is the only available electron acceptor. However, if an increase in OM input in coastal areas causes the SMTZ to shift upwards and become narrower, the active methanogenesis zone may simultaneously expand both in an upward and downward direction. Such expansion of the methanogenesis zone and narrowing of the SMTZ is evident in the Scharendijke basin, as the endogenous methanogenesis rates are equally high in the SMTZ (5-10 cm) and below it at 15-20 cm. Strikingly, methane production was detected with and without added substrates in all tested zones, even in the top 5 cm, and thus above the SMTZ. We note, however, that the sulfate concentrations in our experiment were lower than the in-situ concentrations, which likely enhanced methane production.

Hydrogenotrophic methanogenesis has been traditionally reported as the main methane source in marine sediments, but OM content, salinity and other factors determine which pathways are more prominent (LaRowe et al., 2020; Liu and Whitman, 2008). Our results show that all four main methanogenic pathways were active, as all substrate-amended incubation experiments showed higher rates than the non-amended incubations. At all selected depths, incubations amended with both methanol and H_2_ showed the highest methanogenesis rates. However, the strict H_2_-dependent methylotrophic methanogens Methanofastidiosales and Methanomassiliicoccales were only detected in a narrow zone within the SMTZ in the top sediments and were more abundant from 20 cm downwards. The high abundance of the methylotrophic *Methanolobus* makes it likely that the supplied methanol was used for methane production by both H_2_-dependent and H_2_-independent methylotrophic methanogens, although methanogenesis was seemingly restricted by H_2_ availability (Fig. S1b). Additionally, endogenous fermentation of OM could have supplied CO_2_ for hydrogenotrophic methanogenesis in these samples. At high hydrogen partial pressures, hydrogenotrophic methanogenesis is more thermodynamically favourable than H_2_-dependent methylotrophic methanogenesis (Feldewert et al., 2020). Thus, it is challenging to determine which pathway(s) is/are responsible for the high rates in the MeOH/H_2_ incubations.

Curiously, the second highest rate in the SMTZ was measured in the presence of acetate, although the microbial community composition showed that acetoclastic methanogens were outnumbered by methylotrophic and hydrogenotrophic methanogens. Acetoclastic *Methanosaeta* was observed at much higher relative abundance at 25-35 cm which can probably be explained by lower total cell numbers in deeper sediments (Böer et al., 2009; Qiao et al., 2018). *Methanosarcina* prefers high concentrations of acetate and was probably activated after a long lag phase when no other substrates or competitive micro-organisms were active (Jetten et al., 1992). Methanol-dependent rates were higher than CO_2_/H_2_-dependent rates at all depths, although hydrogenotrophic methanogens from the family *Methanomicrobiaceae* such as *Methanogenium* outnumbered *Methanolobus* and *Methanococcoides* below 10 cm. Therefore, our results suggest that methylotrophic methanogenesis plays an important role in Lake Grevelingen sediments and could even be the main methane source at certain depths. This supports recent studies in various coastal systems that suggest that conversion of methylated substrates might account for a substantial share of methane in marine and coastal sediments (e.g. Zhuang et al., 2016; Tsola et al., 2021; Xiao et al., 2018; Tsola et al., 2024).

### SMTZ hosts a poor methane filter inhabited by ANME archaea and SRB

S-AOM is the major pathway for methane removal in marine sediments, and its occurrence at the Scharendijke basin has been demonstrated before (Bhattarai et al., 2017; Cassarini et al., 2019). To elaborate on earlier findings, we tested for S-AOM potential across the sediment profile, but S-AOM was only observed at two sampling depths, in the SMTZ and just below (10-20 cm). Curiously, no sulfate-dependent methane oxidation signal was detected at 5-10 cm depth, although ANME-2a/b amplicon reads were detected in this layer at 2.4-5% relative abundance (8-10 cm depth) and a small signal of endogenous methane oxidation was detected without added sulfate. The ANMEs might have been inhibited by sulfide in our incubations, as the added sulfate was probably rapidly reduced to sulfide by the highly abundant SRB in the top 10 cm of the sediment. Sulfide toxicity has previously been suggested to inhibit ANME activity in Stockholm archipelago sediments (Dalcin Martins et al., 2024). Sulfide toxicity within the SMTZ might also explain the small niche in which both ANMEs were found, suggesting that high mineralization rates by SRB may further negatively affect the methane filter in coastal sediments in eutrophic sediments. Moreover, ANME archaea are notoriously slow-growing with doubling times varying ∼2-7 months even in laboratory conditions (Meulepas et al., 2009; Nauhaus et al., 2007), with in-situ rates estimated to ∼130 days with reactive transport model (Lenstra et al., 2023). Thus, the lack of appreciable S-AOM could also be associated with the short residence time and low growth rate of the ANMEs in the SMTZ, as suggested by Egger et al. (2016). Alternatively, high rates of organic matter degradation may provide a surplus of H_2_ that is not used up by SRB, which would make S-AOM by reverse hydrogenotrophic methanogenesis thermodynamically unfavourable (Coon et al., 2023). However, most studies based on (meta)genomic characterization support S-AOM driven by direct interspecies electron transfer to their syntrophic partner (Chadwick et al., 2022; McGlynn, 2017). This suggests that both biomass-limitation, due to slow growth, and high sulfide concentrations, could explain the inefficient methane filter in Lake Grevelingen sediments and in similar settings such as the sulfide-rich sediments of Cape Lookout Bight (Coon et al., 2023).

Bhattarai et al. (2017) suggested ANME-3 to be the dominant ANME clade in Scharendijke basin sediments, but our high-resolution 16S rRNA gene profile showed that they only comprised up to 9% of relative archaeal abundances, whereas ANME-2a/b covered up to 15% of the reads in the SMTZ. However, the 16S rRNA gene profile suggests a niche separation for the two ANME clades; ANME-2a/b peaks at 11 cm depth and ANME-3 at 16 cm. At 10 cm, sulfide was still detected at 0.8 mM, suggesting that ANME-3 might be more sensitive to sulfide and thus inhabits deeper zones where sulfide concentrations are negligible. The bacterial community analysis suggests that *Desulfobulbaceae* (DBB) are the bacterial partners for S-AOM in the SMTZ as they correlate with ANME abundance. Interestingly, this DBB branch of ANME-SRB syntrophic partners is usually associated exclusively with ANME-3 (Lösekann et al., 2007). ANME-2a/b is commonly detected in aggregates with the SEEP-SRB1 clade, part of the *Desulfosarcinaceae* family (Murali et al., 2023; Schreiber et al., 2010). *Desulfosarcinaceae* reads were found consistently at ∼2-5% relative abundance throughout the sediment, and genus level classification revealed that from 12-19 cm SEEP-SRB1 reads comprised more than half of the total *Desulforsarcinaceae* reads, thus suggesting that they also had a role in S-AOM, potentially with ANME-2a/b.

To understand all aspects of the sedimentary methane cycle, we looked at the presence of aerobic methane oxidizing bacteria (MOB). Gammaproteobacterial *Methylomonadaceae* reads were detected around the SMTZ, but were likely not active in the sediment. Although MOB have several surviving strategies under oxygen limitation (Reis et al., 2024; Schorn et al., 2024), at the time of sampling, water column ^13^δCH_4_ profiles and potential methane removal rates did not indicate MOB-mediated methane removal in the anoxic bottom water (Venetz et al., 2023, Zygadlowska et al., 2023). Thus, the detected *Methylomonadaceae* probably represent buried cells from the water column MOB community, and a significant contribution of MOB to methane removal in the anoxic sediment is therefore unlikely.

The methane oxidation activity together with the 16S rRNA amplicon profile show that the methane oxidation potential is low in the sediments of the Scharendijke basin, and that methanotrophs are vastly outnumbered by methanogens, explaining the high methane emissions to the water and atmosphere observed at this site (Venetz et al., 2024; Żygadłowska et al., 2024b).

## Conclusion

We investigated the microbial methane cycling potential in the anoxic sediments of marine Lake Grevelingen. We show with incubation experiments of both methane oxidation and production potential combined with 16S rRNA amplicon sequencing analysis that the microbial methane cycle in the sediment is favoured towards methanogenesis, making this system a source of atmospheric methane. The top 60 cm of the sediment harbour a highly diverse archaeal methanogen community, which is distributed along specific niches based on substrate availability. Their metabolic diversity is reflected in the high methanogenic rates from all methanogenic pathways, even in and above the SMTZ. In contrast, methane removal is restricted to the narrow SMTZ. There, ANME archaea are only found in a small niche in the SMTZ, where they oxidize methane via S-AOM in syntrophy with *Desulfobulbaceae*. Our results indicate that anthropogenic eutrophication of coastal waters has the potential to increase methane emissions from these systems, as the changes favour methanogenesis over methanotrophy. Future research is needed to study the methane signature in these systems to understand the contribution of different methanogenic pathways in coastal sediments to the methane emissions.

## Acknowledgement

This research was funded by NESSC NWO-OCW 024002001, SIAM NWO-OCW 024002002 and ERC MARIX 854088. WKL and PDM acknowledge funding by the Dutch Research Council (NWO VI.Veni.222.332 and VI.Veni.212.040, respectively).

