## Supplemental data for "A ubiquitous and diverse methanogenic community drives microbial methane cycling in eutrophic coastal sediments"

### Supplementary figures

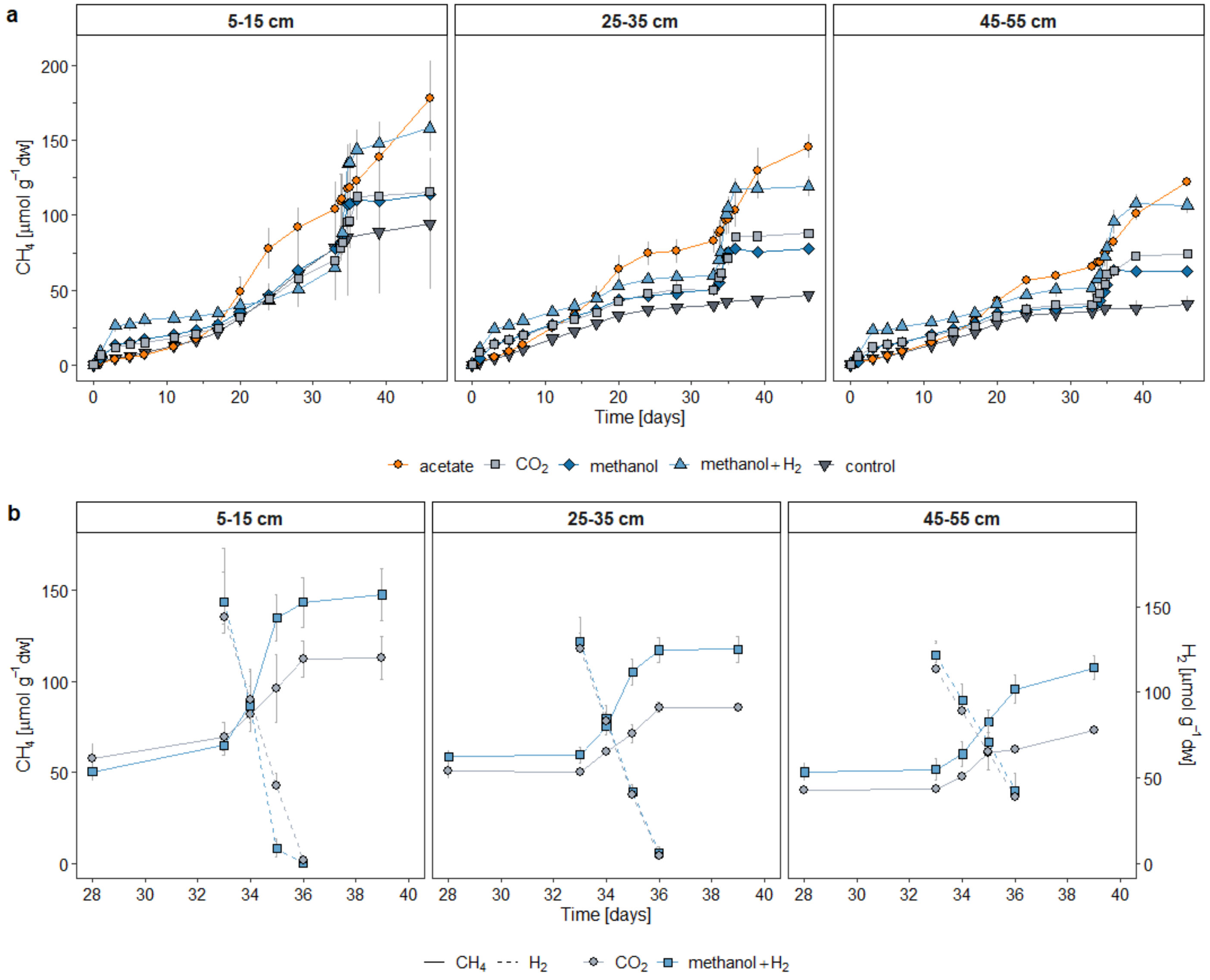


**Fig. S1.** Total methane production in substrate amended and control samples over the whole experiment (a) and during the second substrate amendment where $H_2$ consumption was also measured (b). Substrates were added on day 0 and 33.

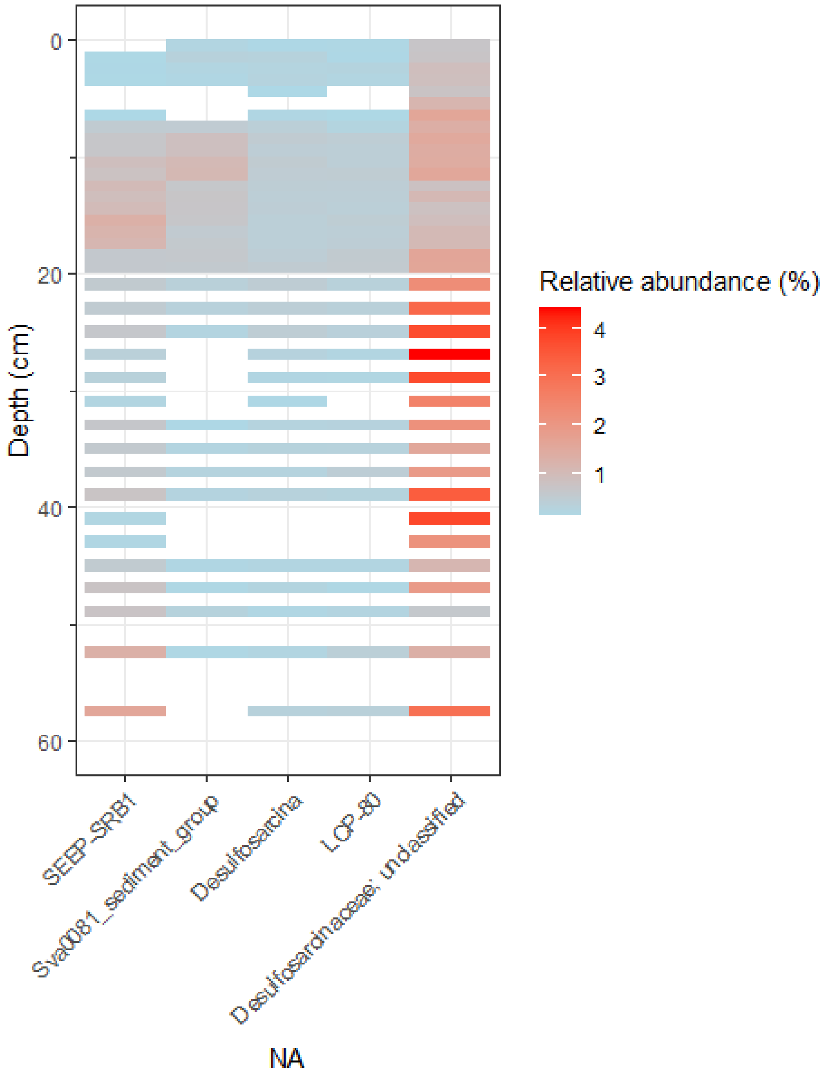


**Fig. S2.** Relative abundance of *Desulfosarcinaceae* reads at genus level.
